# A Clade-D Auxin Response Factor is a Major Regulator of Auxin Signaling in *Physcomitrium patens*

**DOI:** 10.1101/2022.10.25.513742

**Authors:** Carlisle Bascom, Michael Prigge, Whitnie Szutu, Alexis Bantle, Sophie Irmak, Daniella Tu, Mark Estelle

## Abstract

Auxin Response Factors (ARFs) are a family of transcription factors that are responsible for regulating gene expression in response to changes in auxin level. The analysis of ARF sequence and activity indicates that there are two major groups-activators and repressors. One clade of ARFs, clade-D, is sister to clade-A activating ARFs, but are unique in that they lack a DNA binding domain. Clade-D ARFs are present in lycophytes and bryophytes but absent in pteridophytes and spermatophytes. The transcriptional activity of clade-D ARFs, as well as how they regulate gene expression, is not well understood. Here, we report that clade-D ARFs are transcriptional activators in the model bryophyte *P. patens* and have a major role in the development of this species. Δ*arfd1,d2* protonemata exhibit a delay in filament branching, as well as a delay in a key cell differentiation event. Additionally, leafy gametophore development in Δ*arfd1,d2* lines lag behind wild-type. We present evidence that ARFd1 interacts with activating, but not repressing, ARFs. An ARFd1 hypomorph, *arfd1^T653L^*, cannot multimerize. Therefore, we propose a model by which clade-D ARFs enhance gene expression by oligomerizing to DNA-bound archetypal ARFs.

## Introduction

The phytohormone auxin is a key regulator of many developmental processes in plants.

Auxin is perceived within the nucleus by a co-receptor complex consisting of a TRANSPORT INHIBITOR RESISTANT 1/AUXIN F-BOX (TIR1/AFB) protein, a subunit of an SCF E3 ubiquitin ligase complex, and an AUXIN/INDOLE-3-ACETIC ACID (Aux/IAA) protein, a transcriptional repressor. Auxin promotes the TIR1/AFB and Aux/IAA interaction resulting in Aux/IAA ubiquitination and degradation (reviewed in (Leyser 2018)). Once the Aux/IAA proteins are degraded, transcription factors called AUXIN RESPONSE FACTORS (ARFs) activate transcription of auxin-responsive genes (Boer et al., 2014; Tiwari et al., 2003).

ARFs are classed as transcriptional activators or repressors based on either sequence homology or experimental evidence (Tiwari et al., 2003; Finet et al., 2013). Most ARFs have three domains or regions. Near the N-terminus is a B3 DNA binding domain (DBD). This domain promotes ARF binding to the DNA sequence TGTCTC, with wobble in the last two bases (Boer et al., 2014). Adjacent to the DBD lies a large intrinsically disordered Middle Region (MR) that is generally regarded as conferring transcriptional activation to activating ARFs (Tiwari et al., 2003). Finally, archetypal ARFs have a C-terminal Phox/Bem1 (PB1) domain. PB1 domains facilitate dimerization with Aux/IAA repressors as well as other ARFs (reviewed in (Leyser 2018)). The PB1 domains of Aux/IAAs and ARFs have conserved charged regions that drive interactions in a directional manner (Powers et al., 2019; Korasick et al., 2014).

Several publications have focused on the conserved elements of auxin signaling in bryophytes (Thelander et al., 2018). Much of the auxin signaling molecular architecture is conserved in the model moss *Physcomitrium patens* (Prigge and Bezanilla, 2010; Prigge et al., 2010). Auxin is involved in several developmental stages including reproductive organ formation (Landberg et al., 2020), phyllid expansion (Bennett et al., 2014), and colonization by tip-growing cells (Plavskin et al., 2016). Haploid *P. patens* spores germinate to form filaments comprised of cells called protonemata. Protonemal filaments lengthen via a tip-growing apical cell. Early in development, the apical cell is slow growing and chloroplast-rich, generating short cells with perpendicular cross walls as it divides. These cells are referred to as chloronemata. Within days, the apical cell transitions to a fast growing, chloroplast-poor caulonemal cell. These cells are longer than chloronemal cells, and their cell walls are at an oblique angle to the axis of growth. The chloronema-caulonema transition is promoted by the auxin indole acetic acid (IAA) based on work in suspension cell culture with another moss, *Funaria hygrometrica* (Johri and Desai, 1973). Asymmetrical cell divisions off caulonemal cells give rise to gametophores, upon which gametes are exchanged. Therefore, the chloronema-caulonema transition is a necessary step in the *P. patens* life cycle.

Studies of the auxin co-receptor AFB proteins as well as the transcriptional repressor Aux/IAAs provide further evidence for auxin’s role in *P. patens* development. RNAi knock-downs of *PpAFBs* results in plants with striking growth phenotypes including the lack of recognizable caulonemata (Prigge et al., 2010). Similarly, gain-of-function mutations that stabilize Aux/IAAs result in plants with repressed auxin signaling. Developmentally, plants expressing stabilized Aux/IAAs exhibit a delay in the chloronemata-to-caulonemata transition, resulting in an inability to make gametophores (Prigge et al., 2010). Interestingly, *P. patens* plants lacking *Aux/IAA* transcriptional repressors, thereby constitutively responding to auxin, make protonemata that could not be identified as either chloronemata or caulonemata (Lavy et al., 2016). This suggests that repression of some auxin-response genes is required to generate both chloronemata and caulonemata. In total, these studies highlight the importance of auxin to the chloronema-caulonema transition specifically and moss development generally.

The role of individual ARF proteins in bryophyte growth and development is an active area of study. The liverwort *Marchantia polymorpha* has a single activating ARF (MpARF1), a single repressing ARF (MpARF2), and one clade-C ARF (MpARF3) that acts as an auxin-independent transcription factor (Kato et al., 2020; Flores-Sandoval et al., 2018). MpARF1, 2 and 3 each have a single DBD, MR, and PB1 domain. In addition to these proteins, *M. polymorpha* encodes a non-canonical ARF called *MpncARF* that lacks a DBD.(Mutte et al., 2018). Despite the absence of a DBD, *ncarf* lines are partially 2,4-D resistant. Several auxin-sensitive genes are misregulated in the mutant, indicating that ncARFs play an important role in the auxin signaling pathway (Mutte et al., 2018).

In contrast to *M. polymorpha* which has one ARF from each clade, the *P. patens* genome encodes seven clade-A, four clade B, three clade C, and two clade D ARFs (Rensing et al., 2008; Lavy et al., 2016). Similar to the noncanonical ARF in *M. polymorpha*, clade-D ARFs in *P. patens* do not have DBDs. Phylogenetically, D ARFs are in a sister clade to activating A ARFs (Lavy et al., 2016; Mutte et al., 2018; Flores-Sandoval et al., 2018), suggesting that they may function as activators. However, their mode of action and their role in *P. patens* development is unknown.

Here we show that the PpARFds, play a significant role in the *P. patens* auxin response and development. Further, we present evidence that multimerization of the ARFds is key to their function. Lastly, we show that an endogenous auxin-signaling in protonemata plays a role in filament branching, and that the PpARFd’s are required for proper branching.

## Results

### Screen for auxin-resistant mutants identifies ARFs with a key role in auxin signaling

To identify novel components of the *P. patens* auxin-signaling pathway, we screened for mutants resistant to the synthetic auxin, 1-naphthaleneacetic acid (NAA). Seven-day-old tissue regenerated from a homogenized strain of *Physcomitrium patens* harboring the *DR5:DsRed2* auxin-responsive reporter gene (Lavy et al., 2016) was mutagenized using UV light. This tissue was then transferred to medium supplemented with NAA, and visually screened for NAA-resistant (*nar*) phenotypes. Wild-type plants produces rhizoid-like cells on this medium, whereas *nar* mutants produce green protonemal tissue or leafy gametophores (Figure 1A). The activity of the *DR5* reporter was also used to assess auxin response. Fifty-seven mutants were isolated including eighteen new alleles of previously identified *NAR* loci: *DIAGEOTROPICA* (3 alleles), *IAA1a* (6), *IAA1b* (4), and *IAA2* (5) (Lavy and Estelle, 2016; Prigge et al., 2010). Twelve of the remaining thirty-nine mutants with the strongest growth and *DR5* expression phenotypes were selected for whole-genome sequencing. The sequenced genomes were scanned for polymorphisms in auxin- or rhizoid-related genes and for genes with independent mutations in multiple mutants.

**Figure 1.**
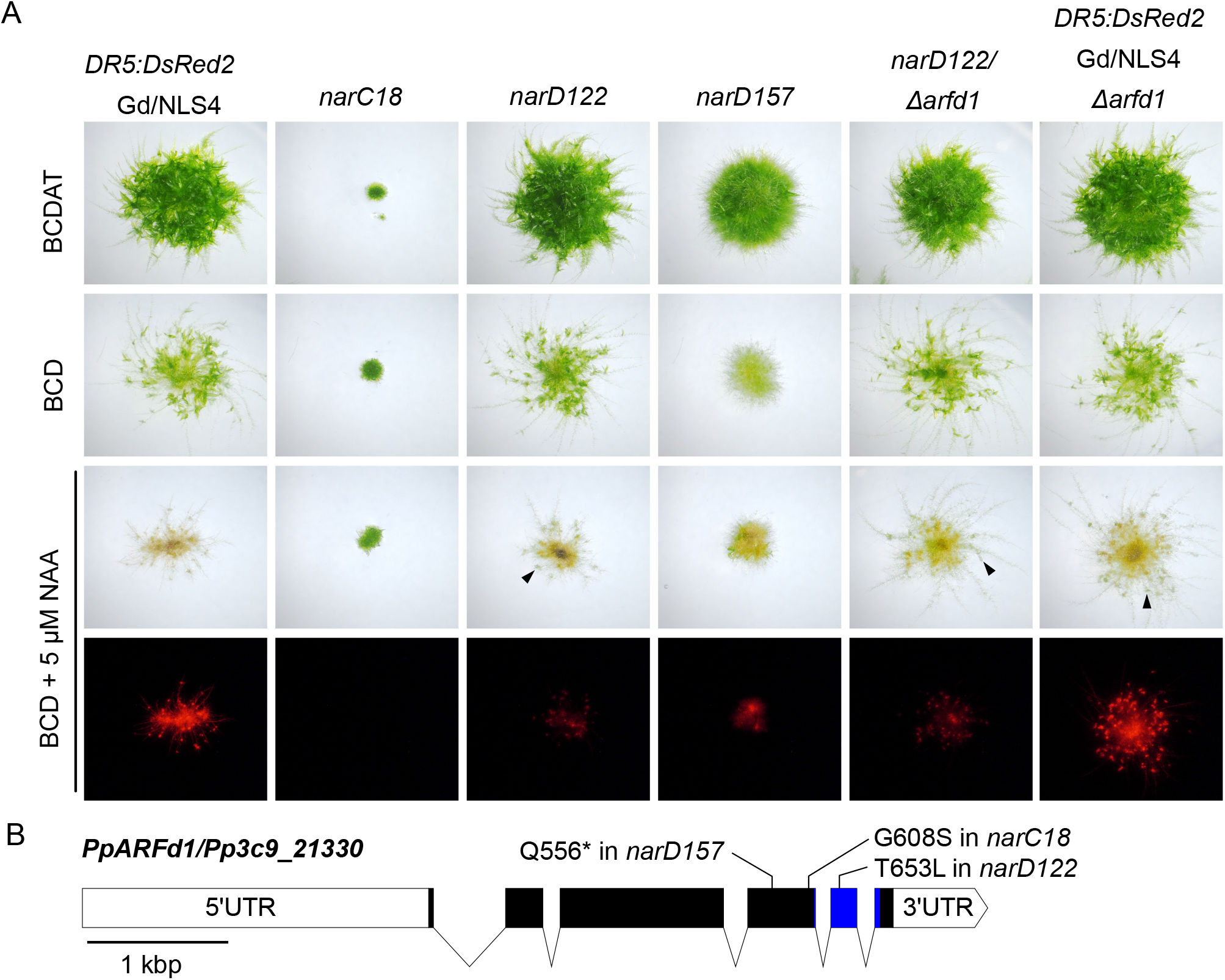
Isolated NAA-Resistant (nar) mutants and causal gene knock-outs. A) *narC18, narD122,* and *narD157* were isolated from irradiated *DR5::DsRed2/*GdNLS4 tissue. *narD122* has a T653L mutation in *ARFd1*. Knocking out *ARFd1* in *narD122* background maintains auxin resistance, suggesting *arfd1^T653L^* is not a dominant mutation. Auxin resistance can be conferred to the *DR5::DsRed2/*GdNLS4 line when *ARFd1* is knocked out. Both *narD122* and Δ*arfd1* genotypes produce leafy phyllids on BCD + 5 μM NAA (black arrowheads). B) Locus of *PpARFd1*, indicating the recovered alleles. UTR-Untranslated Region. Blue region corresponds to PB1 domain.

The *narD122*, *narD157*, and *narC18* mutants all had lesions in the *ARFd1/Pp3c9_21330* locus encoding a clade-D ARF (Figure 1B). *narD157* also had mutations in *ARFa8/Pp3c1_40270*, *PINc/Pp3c10_24880*, and *RSL7/ Pp3c3_8460* while *narC18* a mutation in *ARFb4/Pp3c6_*2137. Because *narC18* and *narD157* contain mutations in addition to those in *PpARFd1,* we elected to focus on *narD122.* To determine if the phenotype observed in *narD122* is due to the mutations in *ARFd1,* we deleted the gene in wild type (*DR5:DsRed*) and in the *narD122* mutant using CRISPR/Cas9 (Figure 1A). Lines with large deletions of *ARFd1* were identified in both backgrounds (Supplemental Figure 1A). Whereas wild-type plants produce rhizoid-like filaments in place of phyllids on leafy gametophores when grown on medium containing 5 μM NAA, *narD122, narD122/*Δ*arfd1*, and Δ*arfd1* plants produced some green phyllids indicating a reduction in auxin response. These results confirm that the auxin response phenotype in *narD122* is due to a defect in *ARFd1*.

### Clade-D ARFs are required for a robust auxin response

There are two ARFd genes in *P.* patens, *ARF1* and *ARFd2* (*Pp3c15_9710*). To determine the contribution of *ARFd2* to auxin response, we used CRISPR/Cas9 to delete *ARFd2* in the wild-type and *arfd1* background, both carrying *DR5::DsRed*. We recovered single *arfd1* and *arfd2* mutants as well as the double knockout (Figure 2, Supplemental Figure 1A). In this experiment, a Gransden ecotype line carrying *DR5::DsRed* (Lavy et al., 2016) had been backcrossed to the Reute ecotype (Hiss et al., 2017) three times. For the remainder of the manuscript, this *DR5* line is the only one used. Where appropriate, comparisons are made using both this line and the Reute wild type.

**Figure 2.**
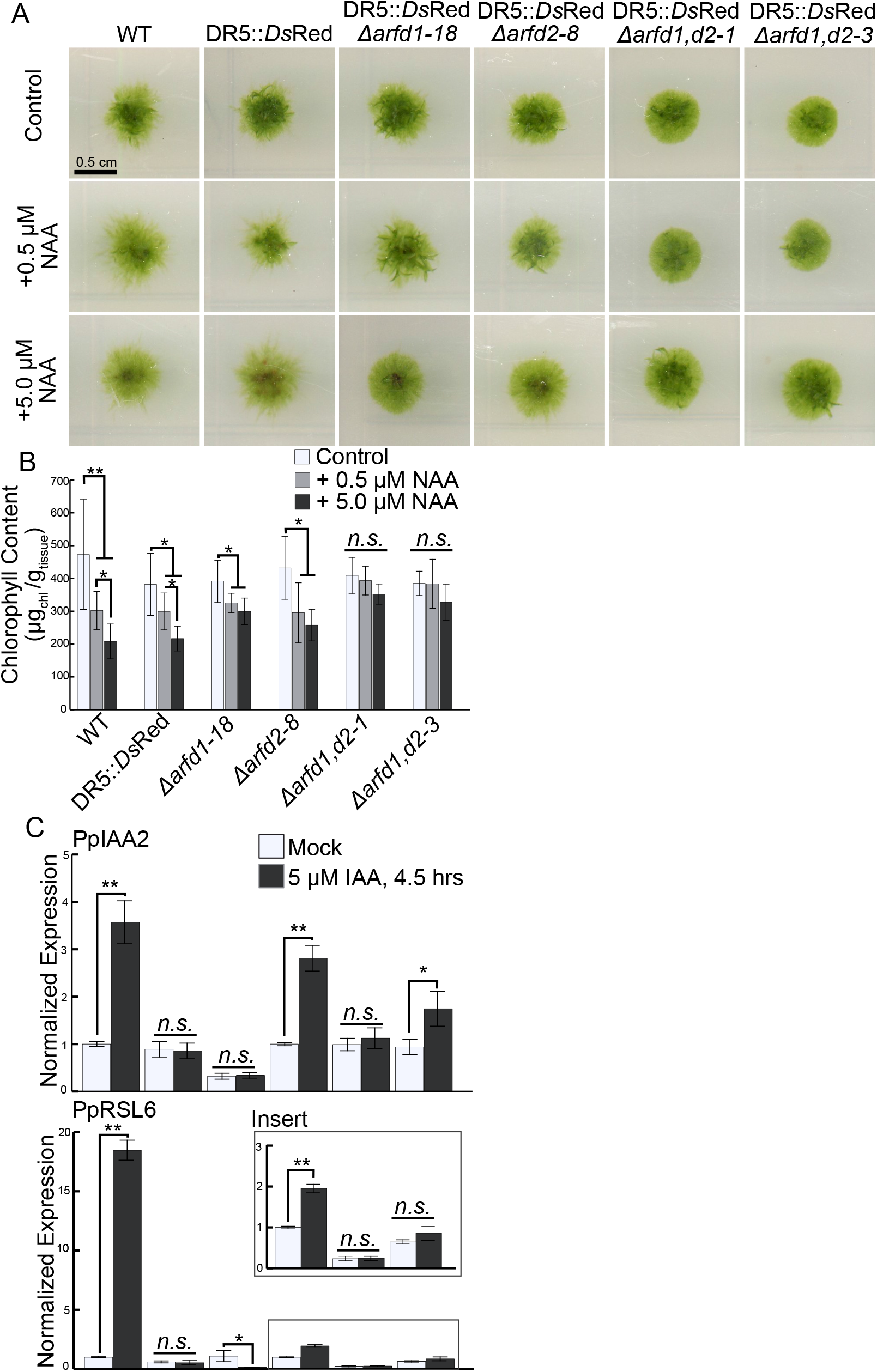
Clade-D ARF knock outs result in changes in morphology and auxin resistance. A) Images of 14-day-old colonies grown from inocula on indicated media. B) Chlorophyll content of 14-day-old colonies grown on indicated media. Average of twelve colonies. Error bars are standard deviation. C) qPCR data showing auxin-induced changes in gene expression for two up-regulated genes (PpIAA2, PpRSL6). Error bars are SEM. *=p-value <0.05, **=p-value <0.001, n.s.=not significant.

Previous reports demonstrated that plants lacking Aux/IAA transcriptional repressors, and are thus constitutively executing the auxin response, have less chlorophyll than wild type (Lavy et al., 2016). Therefore, to assess the level of auxin resistance in the clade-D ARF mutants, we measured the chlorophyll content of plants grown on media with or without NAA (Figure 2). Because of the fourteen-day duration of the growth assay, we used NAA to avoid the photodegradation that can occur with IAA. Nitrogen levels in the medium can also affect chlorophyll levels (Supplemental Figure 1 B,C). Ammonium-deficient (BCD) medium with NAA supplementation has an additive effect in reducing the chlorophyll content. Chlorophyll content in wild-type and *DR5* lines on BCD plus NAA is at or near the detection limit, making comparisons difficult to interpret. Therefore, to accurately measure chlorophyll content, we spot-inoculated moss tissue onto ammonium-supplemented BCDAT medium with or without NAA (Figure 2). After two weeks on BCDAT + 0.5 μM NAA, Reute wild-type plants had 36% less chlorophyll than plants grown without NAA. Similarly, the *DR5* wildtype plants had 22% less chlorophyll on 0.5 μM NAA (Figure 2B). On 5 μM NAA, the two wild-type lines had a 56% and 43% reduction, respectively, compared to the no auxin control. In contrast, each Δ*arfd1,d2* line had only a minor decrease in chlorophyll content when grown on NAA (Figure 2B). Both single mutants had an intermediate response to NAA, suggesting both clade-D ARFs contribute to auxin signaling. Δ*arfd1* plants on 5 μM NAA have 17% less chlorophyll content compared to no NAA. Similarly, Δ*arfd2* plants grown on 5 μM NAA have 32% less chlorophyll than the mock treated counterparts. Δ*arfd1* plants are more resistant than Δ*arfd2* to the effects of exogenous NAA (*p*<0.001, Student’s T-test). This difference suggests that, of the clade-D ARFs, ARFd1 is the main driver of the auxin response. Therefore, we focused subsequent experiments on *ARFd1*.

The lack of a physiological response to exogenous NAA in Δ*arfd1,d2* plants is consistent with ARFds functioning as activators. To obtain more direct evidence for this hypothesis, we analyzed the expression of auxin-regulated genes in the Δ*arfd1,d2* line using qRT-PCR. For comparison, we used two lines with diminished auxin signaling- a IAA2-stablized line (*iaa2-mDII*) and a line lacking all four AUXIN F-BOX genes, (*afb1,2,3,4*) (Supplemental Figure 2). We treated each line with 5 μM IAA (or mock) for 4.5 hours. Finally, we normalized changes in gene expression to the parental line for each mutant (Figure 2C). Neither of the two auxin-induced genes, *IAA2* and *RSL6*, were up-regulated in the *iaa2-mDII* or *afb1,2,3,4* line and there was only nominal up- regulation Δ*arfd1,d2* lines. Taken together, both the physiological and gene expression data point to clade-D ARFs functioning predominately as activators in *P. patens*.

### The PB1 domain is required for ARFd1 function

Our results indicate that the clade-D ARFs are positive regulators of auxin signaling despite the fact that they lack a DBD and presumably cannot bind DNA directly. To gain insight into how the ARFds regulate transcription, we took advantage of the *arfd1^T653L^* mutant identified in the *narD122* line (Figure 1). To confirm that the *arfd1^T653L^* substitution confers an auxin-resistant phenotype, we employed CRISPR/Cas9 in combination with a oligonucleotide carrying the mutation to recreate the T653L substitution in an ARFd1-mYPet/Δ*arfd2/*DR5::DsRed background. Similarly, we generated lines containing a Y643* mutation, which generateds a premature stop codon just before the PB1 domain (Supplemental Figure 3A,B). We determined the level of auxin resistance in these lines by measuring the chlorophyll content of tissue grown on medium with or without 5 μM NAA for twenty-one days (Figure 3C, D). Tissue of the control (*DR5)* and ARFd1/Δ*arfd2* lines had 2.62 and 2.49-fold, respectively, more chlorophyll on mock medium compared to NAA-supplemented medium. Similar to results presented in Figure 2, chlorophyll levels in the Δ*arfd1,d2* lines were not affected by NAA treatment. Likewise, independent Y643* lines demonstrated a similar level of NAA resistance indicating that the PB1 domain is necessary for ARFd1 function. Independent *arfd1^T653L^* lines exhibit an auxin sensitivity that is intermediate between the control and Δ*arfd1,d2* lines. Therefore, we conclude that arfd1^T653L^ retains some activity.

**Figure 3.**
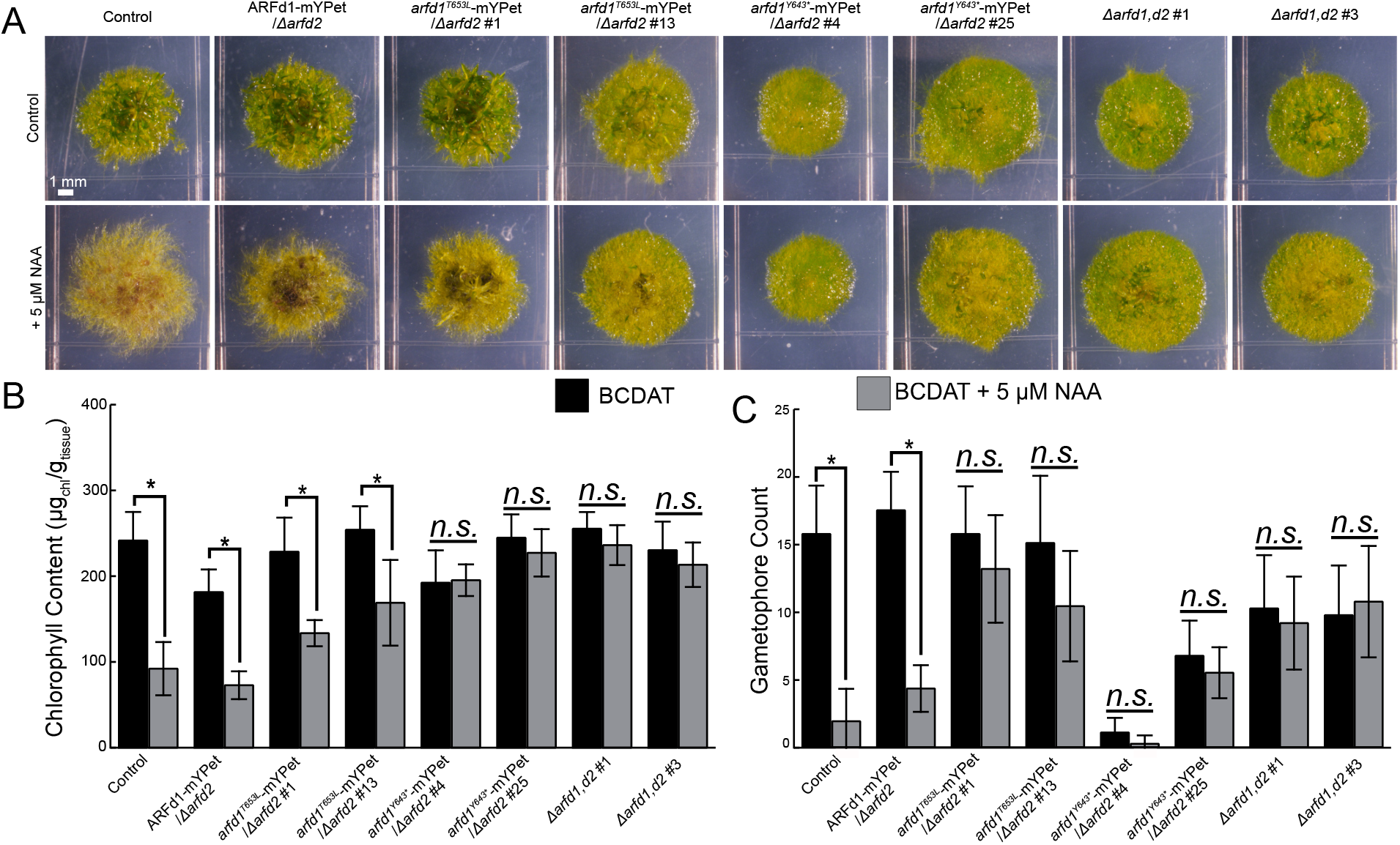
ARFd1 function depends on PB1 domain. A) 21-day-old colonies of ARFd1 mutants grown +/− 5 μM NAA. Control line is *DR5::DsRed* parental line. B) Chlorophyll content of whole colonies imaged (n=6 for each line and treatment). *=*p<0.*001. C) Average gametophore count for n=6 colonies grown +/− 5 μM NAA for 21 days.

In addition to affecting chlorophyll levels, 5 μM NAA inhibits gametophore development in wild-type *DR5* and *ARFd1-mYPet* lines. While wild-type plants initiate gametophore-like structures on 5 μM NAA, these structures develop rhizoids instead of phyllids. In contrast, there are fewer gametophores produced on Δ*arfd1,d2* plants indicating that at least one functional ARFd is required for proper gametophore development. Consistent with auxin resistance, auxin does not affect gametophore number in the double mutant. Again, the *arfd1^T653L^* lines exhibit an intermediate phenotype. They develop gametophores at levels comparable to control plants (Figure 3C) but interestingly gametophore number is not affected by NAA treatment (Figure 3A,C). These observations are consistent with published findings of auxin-insensitive *Physcomitrium* mutants (Prigge et al., 2010; Plavskin et al., 2016).

### The ARFd proteins have a role in the chloronema-caulonema transition in an ammonium-dependent manner

Previous studies have shown that auxin promotes the chloronema-caulonema transition. Consistent with this, we found that Δ*arfd1,d2* colonies are noticeably round when growing on ammonium-supplemented medium (BCDAT). These observations suggest that the ARFd proteins are required for the development of faster-growing caulonemata. However, on ammonium-deficient medium (BCD), we do not detect a difference in colony circularity (Supplemental Figure 1D). To address this issue directly, we regenerated plants from single protoplasts on both ammonium-supplemented and ammonium-deficient media (Figure 4A) and determined the level of caulonemal development by measuring cell cross-wall angle. The angle of the cross-wall is widely recognized as a key morphological distinction between chloronemata and caulonemata. Chloronemata have cross-walls that are perpendicular to the growth axis of the filament, while caulonemata cross-walls are oblique. The wild-type *DR5* line on BCD had an average cross-wall angle of 114.1±15.9° (Figure 4B). Consistent with the morphological distinction between chloronemata and caulonemata, the cross-wall data form a bimodal distribution around 110°. Therefore, we used 110° as our cutoff to differentiate chloronemata and caulonemata in our data. In *DR5,* 40% of measured cross-walls fell below this threshold and were scored as chloronemata, while 60% of cross-walls belonged to caulonemata (Figure 4B). Interestingly, both Δ*arfd1,d2* mutants have nearly the same cross-wall average (114.1±16.6° and 113.9±16.1°, respectively) and only slight differences in distribution (42% chloronema, 58% caulonemata for each) compared to the parental *DR5* line (Figure 4A,B). Neither T-tests of the averages and Kolmogorov-Smirnov test of the distributions yielded a significant difference. Taken together, these data suggest that the Δ*arfd1,d2* has a normal ability to make caulonemata on ammonium-deficient BCD medium.

**Figure 4.**
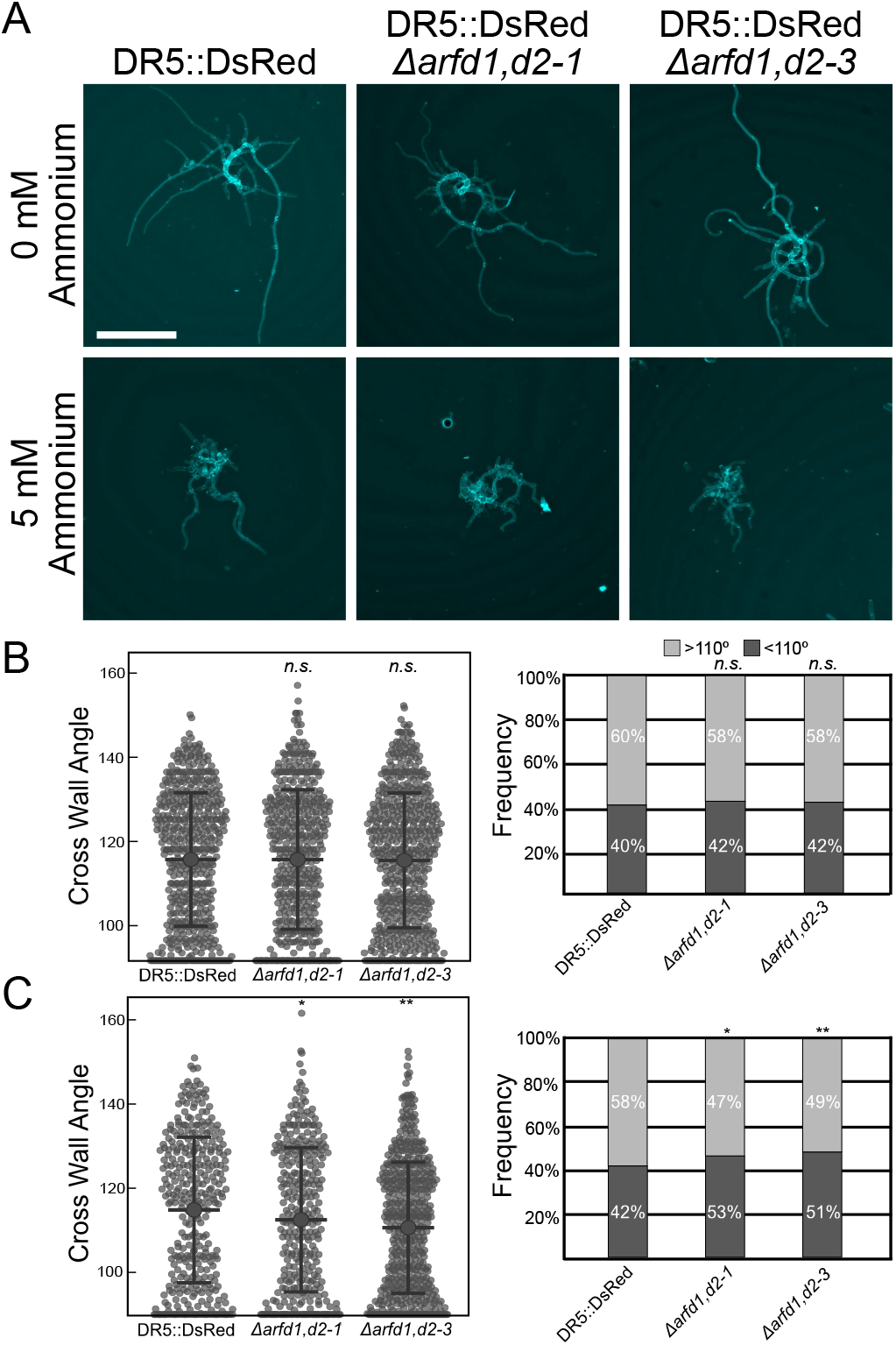
*arfd* lines have an ammonium-dependent caulonemal development delay A) Images of representative seven-day-old plants grown from regenerated protoplasts of indicated line ammonium deficient (above) or ammonium-supplemented (below) medium. Plants stained with Calcofluor for imaging, scale bar is 500 μm. Cross wall measurements of plants grown on ammonium-deficient (B) and ammonium-supplemented medium (C) and frequency of cross walls binned as >110° or <110°. Error bars are standard deviation, *=p<0.05, **=p<0.001.

In contrast, the presence of ammonium in the medium had a significant effect on caulonemal development. In these conditions, the average cross-wall angle for *DR5* plants was 114.8±17.3°, with 42% of measured cells counting as chloronema (Figure 4C). Meanwhile, both Δ*arfd1,d2* lines had a slight, but significant, enrichment in chloronemata. This was true both in the average cell wall angle (112.5±17.1° and 110.6±15.6°, respectively), and percent of cells with cross walls less than 110° (53% and 51%, respectively). Conducting a Kolmogorov-Smirnov test on the distribution of cross-wall data determined that the detected shift was significant.

Given that the chloronemal enrichment was slight in Δ*arfd1,d2,* we hypothesized that ammonium-dependent delay in caulonemal differentiation should be more pronounced in plants with more severe mutations in the auxin signaling pathway. Therefore, we grew *iaa2-mDII* and *afb1,2,3,4* plants from regenerated protoplasts on medium with or without ammonium. Independent of ammonium levels, both *iaa2-mDII* and *afb1,2,3,4* protonemata are enriched in chloronemata (Supplemental Figure 4). These observations are in line with previous reports that auxin signaling is required for caulonemata development (Supplemental Figure 4). For each mutant, the ammonium-dependent chloronemal enrichment was enhanced compared to wild type. In the absence of robust auxin signaling, the contribution of ammonium-sensing to the chloronemal-caulonemal transition is more apparent.

Taken together, auxin signaling and nutrient sensing are drivers of the chloronema-caulonema transition. ARFds are required for caulonemal development, but only in nutrient-rich conditions.

### Clade-D ARFs are required for proper branch filament initiation

Thus far, we have demonstrated that clade-D ARFs are required for a robust auxin response and for caulonemal-development in high-nitrogen conditions. Given that Δ*arfd1* plants have reduced DR5::DsRed expression (Figure 1), we hypothesized that clade-D ARFs function as activators, and therefore may exhibit additional developmental defects when high levels of auxin signaling are required. Foundational work in the moss *Funaria* suggests that IAA regulates the number of filament branches (Johri and Desai, 1973), as *Funaria* protonemata grown in medium supplemented with IAA developed more branches. However, more recent findings employing exogenous NAA, and increasing endogenous auxin in *Physcomitrium* with *pinapinb* mutants, suggests that auxin has little effect on the number of branches per filament (Thelander et al., 2018). In our conditions, we observed increased *DR5* activity in young filament branches (Figure 5A), suggesting that endogenous auxin signaling may play a role in protonemal branching. Therefore, we compared filament branch emergence frequency in the *DR5* and Δ*arfd1,d2* lines. We scored each filament by the position of the first branch relative to the apical cell. Filaments from the *DR5* control line typically branch at the second subapical cell (44.8%) (Figure 5B, D). On rare occasions, the first subapical cell will branch (<5%, Figure 5D). Exposure to 5 μM IAA for three days induces premature branch emergence. Indeed, auxin stimulates branch initiation in the first subapical cell most frequently (45.2%), even occasionally causing filaments to branch at the apical cell (14.0%) (Figure 5C, D). This observation supports a model in which the level of auxin contributes to the timing of branch initiation.

**Figure 5.**
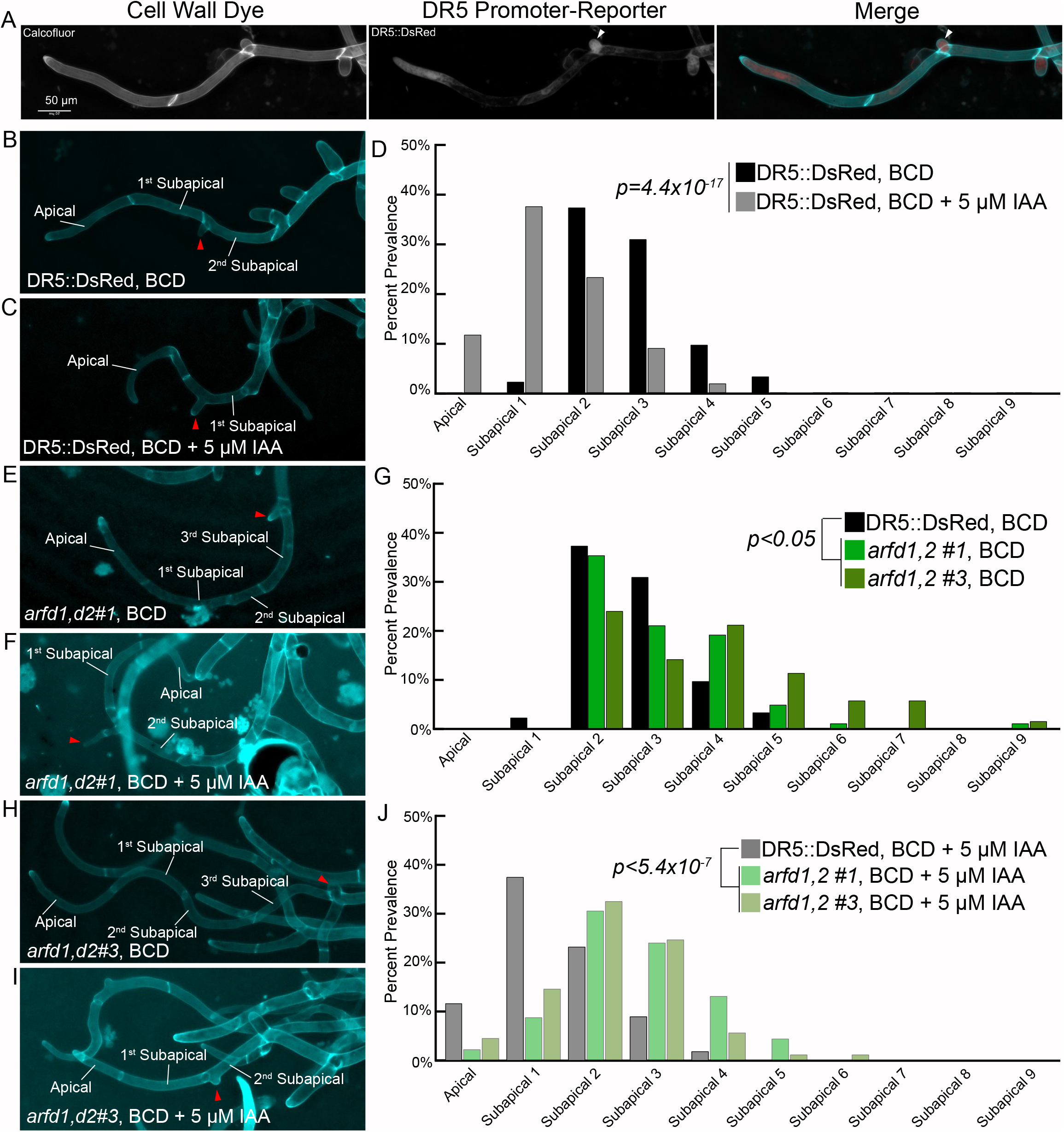
Branch initiation is delayed in *arfd* mutants. A) Representative micrographs from *DR5* line showing that auxin signaling increases in filament branch cells (white arrowhead). B,C) *DR5* line grown on BCD±5 μM IAA, red arrowhead highlights emerging branch. D) Branching in control plants occurs most often in the second subapical cell (black bars), but in the first subapical cell when grown on auxin (grey). E,G,H) Δ*arfd1,d2* filaments have a delay in branching emergence. F,J,I) 5 μM IAA treatment of Δ*arfd1,d2* plants partially rescues branch emergence phenotype. Statistics for branching determined by assigning numerical values to cell number (apical=0, subapical=1, etc) and performing students t-test.

In contrast, both Δ*arfd1,d2* lines exhibit a delay in branch emergence (Figure 5E, H, G). While both mutants frequently branch at the second subapical cell (42.5% and 28%, respectively), there is an increase in the number of late-forming branches (Figure 5G). Indeed, 22-25% of Δ*arfd1,d2* filaments had formed the first branch in their fourth subapical cell, compared to 11.5% in the *DR5* line. Additionally, 3-15% of mutant filaments had branches emerging after the sixth subapical cell, something that was never observed in the *DR5* line. Similar to *DR5* plants, we observed a stimulation of branch emergence in Δ*arfd1,d2* lines when grown on 5 μM IAA (Figure 5F, I, J). On auxin, the frequency of branching at the second subapical cell changed to 36.8% and 39.2% for the double mutant lines, respectively. Additionally, branching on or after the sixth subapical cell occurred in <2% of mutant filaments when grown on auxin. These results show that auxin response is a major factor in determining filament branch timing.

These results indicate that auxin plays a fundamental role in protonemal filament branching. We sought to validate our studies by measuring branch emergence in other auxin signaling mutants. We scored branch emergence in a line expressing a stabilized PpIAA2 (*iaa2-mDII)* and a line lacking the four auxin co-receptor F-Box proteins (*afb1,2,3,4)* encoded by the *P. patens* genome. Both lines have impaired auxin signaling (Figure 2C) and exhibited a delay in filament branching (Supplemental Figure 5), confirming a role for auxin in branch formation.

### ARFd1 is expressed in protonemal filaments and the base of growing gametophores

*PpARFb4* expression is restricted to the distal ends of protonemal filaments (Plavskin 2016). By contrast, activating ARF expression pattern in *Physcomitrium* remains unknown. Therefore, we investigated ARFd1 expression using ARFd1-mYPet signal (Figure 6). Given the role of ARFds in the chloronemal-caulonemal transition, we hypothesized ARFd1 would be enriched in a specific cell type. To this end, measured mYPet signal in seven-day-old *ARFd1-mYPet*/Δ*arfd2/*DR5::*DsRed* plants regenerated from protoplasts (Figure 6A). We then measured ARFd1-mYPet signal and the angle of the proximal cell wall. However, ARFd1-mYPet signal was not significantly different across cell type using the same 110° delineation between chloronemata and caulonemata (Figure 6B).

**Figure 6.**
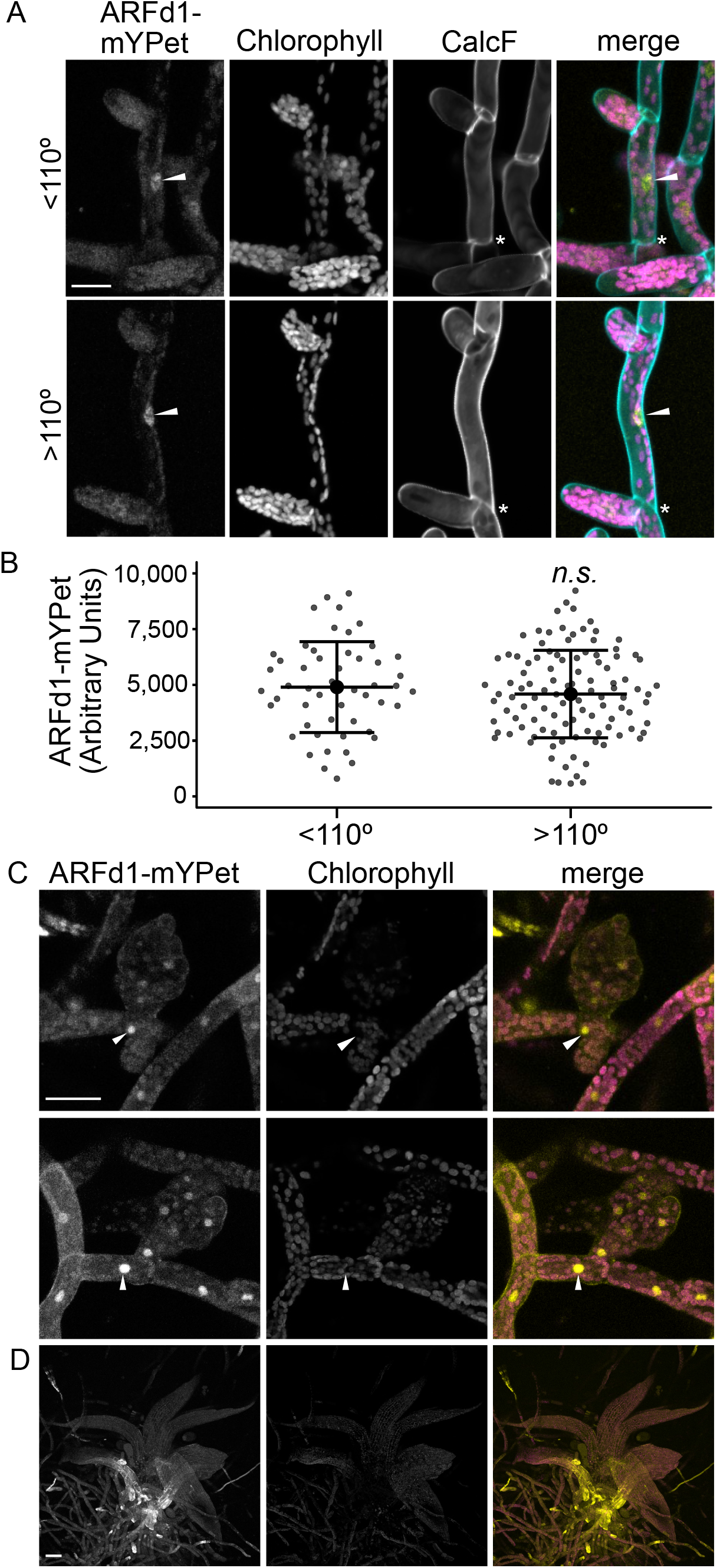
ARFd1 expression pattern A) Representative micrographs from seven-day-old plants regenerated from pARFd1::ARFd1-mYPet/Δarfd2/DR5::DsRed plants. Top is a cell with the proximal cell wall (asterisk) of <110° (chloronema cell), bottom is >110° (caulonema cell). White arrowhead highlights nuclear ARFd1mYPet signal. Scale bar is 25 μm.) Nuclear ARFd1mYPet signal binned by cell wall angle. C) Representative micrographs of young gametophore buds. ARFd1mYPet is enriched in the protonemal cell at the base of the developing gametophore (white arrowheads). Scale bar is 50 μm. D) ARFd1mYPet signal is absent in mature gametophores. Scale bar is 100 μm.

Δ*arfd1,d2* lines have delayed gametophore development (Figure 3C). Therefore, we searched ten-day-old regenerated homogenized *ARFd1-mYPet*/Δ*arfd2/*DR5::*DsRed* tissue for young gametophores (Figure 6C). We consistently detected a bright ARFd1-mYPet signal in the protonemal cells at the base of the developing gamotophore (Figure 6C, white arrowhead). This observation is coherent with clade-D ARFs playing an important role in gametophore initiation. We did not detect ARFd1-mYPet in mature gametophores (Figure 6D), suggesting that ARFd1 is involved in gametophore bud initiation, but not necessarily in their development.

### ARFd1 function requires oligomerization

We next sought to determine a mechanism by which clade-D ARFs can activate gene expression without a DBD. One hypothesis is that clade-D ARFs interact with canonical ARFs bound to DNA to affect gene expression. Most ARFs have a PB1 dimerization domain, and because T653 is in the PB1 domain of ARFd1 (Figure 1B), this mutation could disrupt the ability of ARFd1 to interact with archetypal ARFs. To test this possibility, we conducted a yeast-2-hybrid assay with full-length ARFs to assess their ability to interact with to ARFd1 (Figure 7A). We found that ARFd1 interacts with both full-length ARFa4 and ARFa8. Conversely, there is minimal interaction with ARFb1, a repressing ARF. Curiously, full-length *arfd1^T653L^* also interacted with ARFa4 and ARFa8, suggesting that the mutation does not generally affect ARFd1’s ability to dimerize with other ARFs.

**Figure 7.**
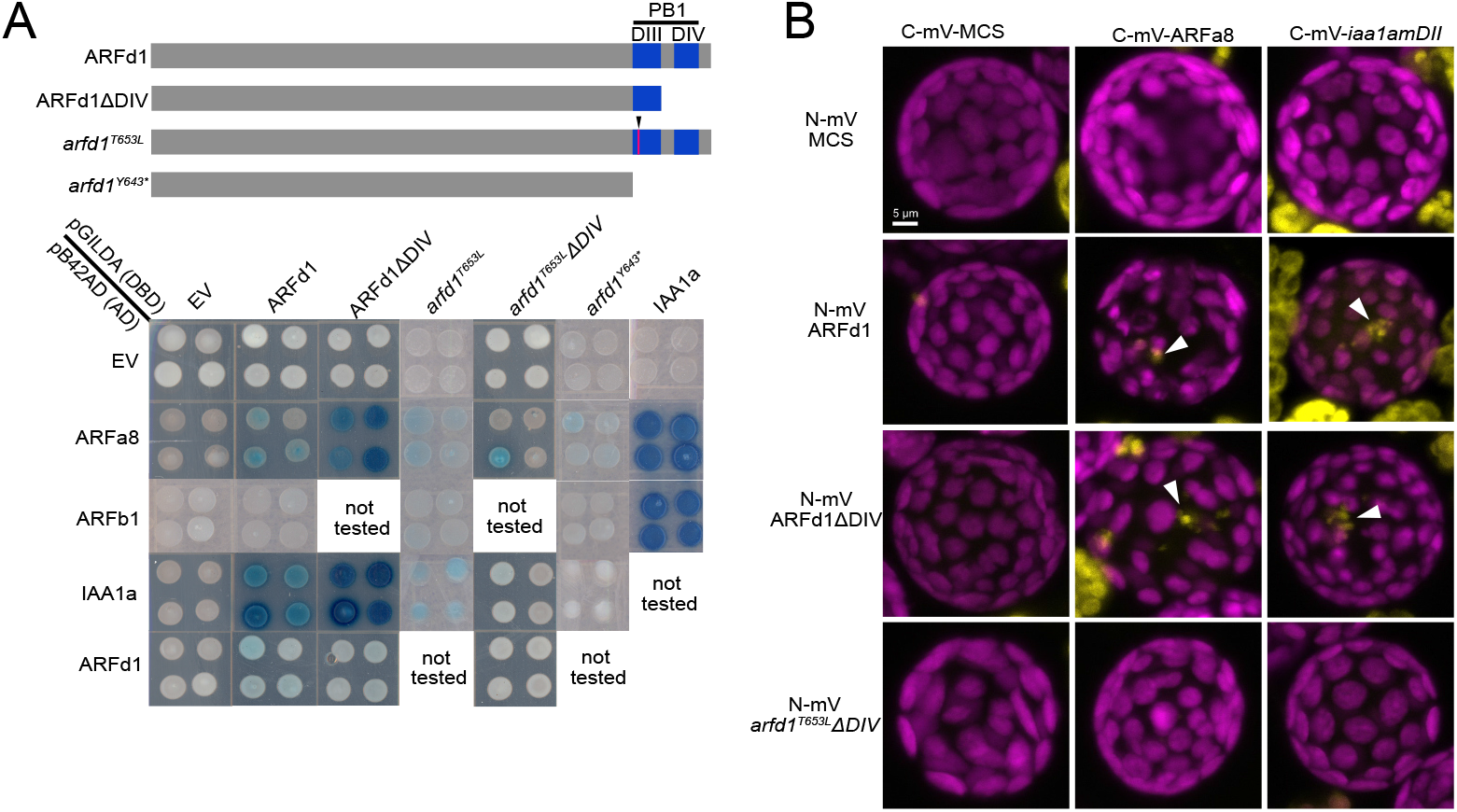
PpARFd1 requires oligiomerization to function. A) A diagram of mutant ARFd1 sequences used, as well as the results of yeast-two-hybrids. Blue indicates positive interaction. ARFd1 interacts with both the activating ARFa8 as well as the transcriptional repressor PpIAA1a. Limiting dimerization through DIII reveals that *arfd1^T653L^* disrupts DIII function. B) Split YFP assay validating ARFd1-IAA1a and ARFd1-ARFa8 interaction is abolished when *arfd1^T653L^* protein can only dimerize through DIII. ARFd1 interaction with both IAA1a and ARFa8 occurs in puncta (white arrowheads).

PB1 domains consist of separate positively and negatively charged surfaces (formerly DIII and DIV respectively) that facilitate oligomerization through surface charge interactions in a head to tail fashion (Korasick et al., 2014; Powers et al., 2019). For ease of discussion, we will refer to the two faces as DIII and DIV. The T653L mutation affects the DIII face of ARFd1 (Figure 7A), and therefore may affect the interaction with the DIV of other ARFs. To test this possibility, we made ΔDIV versions of ARFd1 and *arfd1^T653L^* and performed yeast two hybrid assays. ARFd1ΔDIV retains the ability to interact with ARFa8, but not ARFb1. However, *arfd1^T653L^*Δ*DIV* cannot interact with ARFa8 indicating that it is the wild-type DIV domain that is responsible for the *arfd1^T653L^*-ARFa8 interaction in yeast. We verified this result in plants using split yellow fluorescent protein (YFP) assays in *P. patens* protoplasts. Interestingly, we observed interaction as sub-nuclear puncta (Figure 7B).

We also tested the ability of these proteins to interact with PpIAA1a. While full-length *arfd1^T653L^* interacted with PpIAA1a just as well as full-length ARFd1 in both yeast and in split YFP experiments, *arfd1^T653L^*Δ*DIV* did not (Figure 7A). As a control, we tested the ability of the *arfd1^Y643^\** protein to interact with ARFa8 and IAA1a. As expected, *arfd1^Y643^\** did not interact with either protein. This confirms that ARFd1 interacts with other ARFs and Aux/IAAs solely through the PB1 domain.

Taken together, these results indicate that the T653L mutation disrupts the function of one of the two charged faces in the PB1 domain. The remaining face (DIV) is sufficient to enable interaction with ARFa8 and IAA1a and partial ARFd1 function in plants. The fact that both faces are required for full activity strongly suggests that ARFd oligomerization contributes to its activity. This is the first evidence in any system that ARF oligomerization is important for its function.

## Discussion

Unlike other members of the ARF family of transcription factors, the clade-D ARFs lack a DNA binding domain. Previous work with another bryophyte, *M. polymorpha*, found that loss of the single clade-D ARF results in effects on auxin regulated transcription (Mutte et al., 2018). However, the role clade-D ARFs play in plant development, as well as how they affect gene expression, were open questions. Our experiments demonstrate that clade-D ARFs function primarily as transcriptional activators and have an important role in the progression of caulonema and gametophore development in *P. patens*.

Clade-D ARFs have a clear role in *P. patens* development because Δ*arfd1,d2* colonies have a higher circularity than wild-type. Previous work has associated high colony circularity with decreased auxin signaling. An increase in colony circularity has been interpreted as an inability to make caulonemata (Plavskin et al., 2016; Lavy et al., 2016; Prigge et al., 2010). When we directly tested the ability for Δ*arfd1,d2* to make caulonemata using regenerated protoplasts, we were surprised to find that Δ*arfd1,d2* plants can make wild-type levels of caulonemata on ammonium-deficient BCD medium. However, Δ*arfd1,d2* plants grown on ammonium-supplemented BCDAT medium have a delay in caulonemal development. This suggests that a nutrient-sensing developmental pathway masks the developmental phenotypes of Δ*arfd1,d2.* Interestingly, even strong auxin-signaling mutants show a nutrient-dependent chloronema-caulonema transition. Future work will have to interrogate if the nutrient-dependent pathway cross-talks with the auxin signaling pathway, or if it is truly a parallel genetic pathway in the chloronemal-caulonemal transition.

Independent of ammonium levels, protonemal branching is delayed in Δ*arfd1,d2*. We observed a similar branching delay in strong auxin signaling mutants (*iaa2-mDII, afb1,2,3,4*). These data suggest that protonemal branch formation is at least in part an auxin-driven process, in line with foundational studies (Johri 1973). In support of this, exogenous IAA results in premature branching relative to the apical cell. Yet, recent work using *pina,pinb* mutants and NAA treatment found no connection between auxin and caulonemal branches (Thelander 2018). In our branching experiments, we did not differentiate chloronemata and caulonemata. Additionally, Thelander et al. measured branching as branches per cell, and did not address the timing of branch emergence relative to the apical cell. Taken together, we are confident that the timing of protonemal branching is at least partially informed by levels of auxin signaling. Additionally, clade-D ARFs are a key regulator of the auxin signaling required to branch.

In addition to the delay we observed in protonemal branching, Δ*arfd1,d2* plants have delayed gametophore development. Interestingly, the gametophores that do develop on Δ*arfd1,d2* are phenotypically wildtype. This would suggest that the role clade-D ARFs play in gametophore development is restricted to initiation. In line with these observations, we detected an enrichment of pARFd1::ARFd1mYPet signal in the protonemal cell at the base of early gametophore buds. While Δ*arfd1,d2* plants have an ammonium-dependent delay in the chloronemal-caulonemal transition, we did not detect a significant difference in pARFd1::ARFd1mYPet signal between the two cell types. There was no cell-specific expression difference even when accounting for ammonium levels in the medium (data not shown). In general, the Δ*arfd1,d2* phenotypes are similar to, albeit less severe than, those of the strong *iaa2-mDII* and *afb1,2,3,4* mutants. This leads us to a model whereby clade-D ARFs are involved in general auxin-mediated gene activation events (Figure 8). When and where auxin-mediated gene activation events occur within the plant is determined by some other factor.

**Figure 8).**
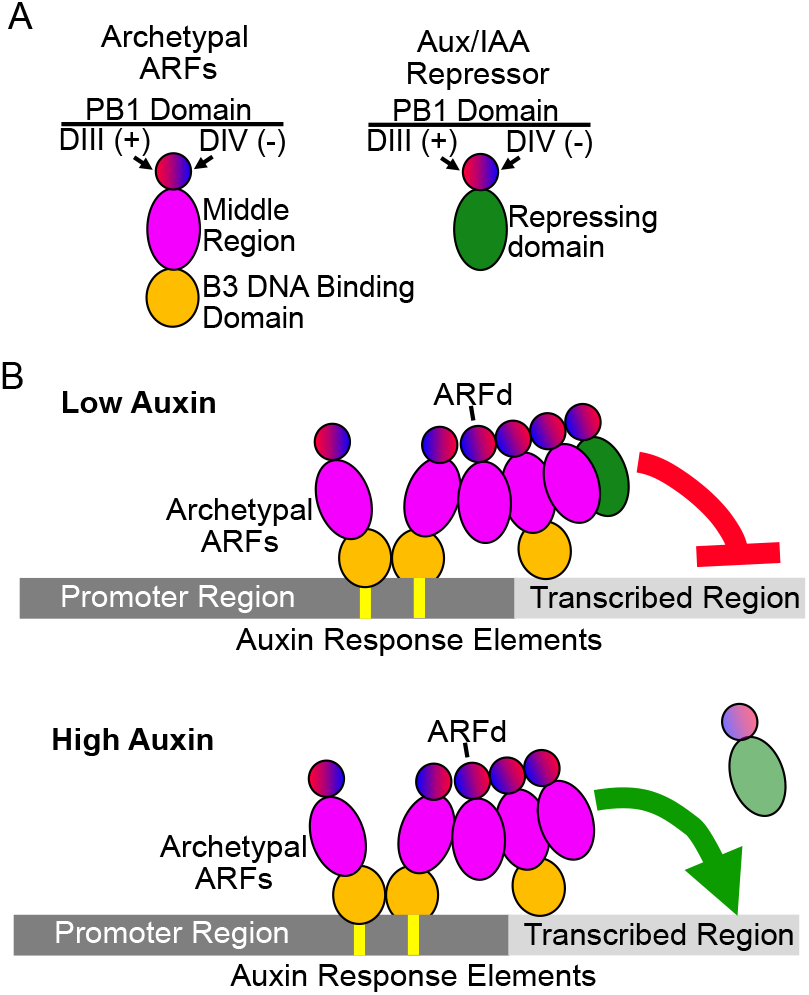
A model for clade-D ARF function A) Schematic of the domains of ARFs and Aux/IAA Repressors B) In low auxin conditions, ARFds facilitate the oligomerization of archetypal ARFs as well as Aux/IAAs to an auxin-inducible gene. Once the Aux/IAA is removed in an auxin-dependent manner, clade-D ARFs activate gene expression in conjunction with archetypal clade-A ARFs.

PpARFd1 can activate gene expression despite lacking a DNA binding domain. Our data demonstrate that an intact PB1 domain is necessary for full function. PB1 domains facilitate interaction between ARFs and Aux/IAA repressors (Vernoux et al., 2011). ARF-ARF interactions can occur via the PB1 domain in oligomerization (Korasick et al., 2014). However, most models suggest that ARF-ARF interaction primarily occurs via the dimerization domain adjacent to the DBD, and only occur when bound to DNA (Boer et al., 2014). Our interaction experiments indicated that PpARFd1 interacts with both activating ARFa8, as well as PpIAA1a, presumably through the common PB1 domain. PB1 domains have two binding surfaces-positive and negative. By limiting protein-protein interactions to the positive face of ARFd1, we demonstrated that the *arfd1^T653L^* mutation results in an ARF that can only dimerize. The most likely model is that clade-D ARFs interact with DNA-bound archetypal activating ARFs (e.g. PpARFa8) and enhance transcriptional activation. *Arfd1^T653L^*/Δ*arfd2* plants have intermediate phenotypes between ARFd1/Δ*arfd2* and Δ*arfd1,d2*, suggesting that ARFd1-mediated transcriptional enhancement can occur through dimerization. However, for wild-type levels of auxin signaling ARFd1 has to multimerize with additional factors. One possible scenario is that ARFd1 recruits additional activating ARFs, either ARFa or ARFd, through their PB1 domains (Figure 8). Our data also indicate that ARFd1 can bind the transcriptional repressor PpIAA1a suggesting that ARFd1 function is auxin regulated via AFB dependent degradation of the Aux/IAAs.

The work presented here demonstrates that clade-D ARFs are a key player in bryophyte auxin signaling. Additionally, we describe that oligomerization is essential to PpARFd1 function. Curiously, clade-D ARFs were lost in pteridophytes and spermatophytes. Future research should investigate if extant ARFs, or other proteins entirely, in vascular plants perform the oligomerization function that clade-D ARFs conduct in bryophytes. Further work will also have to address if oligomerization is unique to clade-D ARFs, or to bryophyte ARFs in general.

## Materials and Methods

### Plant tissue and growth conditions

For all assays, plants were grown in a Percival growth chamber under constant light conditions at 25°C. Transgenic plants were made using DR5::*Ds*Red originally made in the Gransden ecotype backcrossed to the Reute ecotype of *Physcomitrium patens* three times (Hiss et al., 2017). Formulations of BCD(AT) media were used as they were described previously (Cove et al., 2009). All transformations of *P. patens* were performed via the PEG-mediated described previously (Schaefer et al., 1991).

### Mutant screen

The *DR5:DsRed2/NLS-4* strain (*DR5:DsRed/35S:aacC1* inserted into Pp108B between *Pp3c20_980* and *Pp3c20_1020*; *2×35S:NLS-GFP-GUS/35S:nptII* inserted randomly) was sterilely homogenized in water in a Waring blender with an autoclaved MC1 mini container, spread onto BCDAT plates containing 100 mg/l gentamicin and 25 mg/l G418 overlain with 90mm cellophane sheets (AA Packaging), and grown for 1 week. In the course of this work, it was discovered that the gentamicin resistance cassette, *35S:aacC1*, confers resistance to both gentamicin and G418. Plates were then placed in a Stratalinker UV Crosslinker (Stratagene) and, with lids removed, irradiated with doses of 254-nm light. The plates were closed, taped, and stored in the dark for 24 hours to block photoreactivation of pyrimidine dimers. For the UV dose-response evaluation, ten doses ranging from 0 to 500 mJ/cm^2^ were tested before a dose of 150 mJ/cm^2^ was chosen for mutagenesis because it caused a mortality rate of approximately 40%. After UV and dark treatments, the cellophanes were transferred to BCD plates with 15 μM NAA (dose-response treatments), 1 μM NAA (“narB” round), or 5 μM NAA (“narC” and “narD” rounds). 5 μM NAA proved to be the most effective concentration and all mutants were isolated from 12 (narC) and 18 (narD) plates. After a 2-4 weeks, the plates were screened for mutants that produced green protonemata and/or leafy gametophores. A few cells of 224 candidate mutants were spotted onto new BCD plates containing 5 μM NAA and re-evaluated after two weeks. This yielded 57 mutants (with fifteen additional candidates selected later based on reduced *DR5* expression). The initial 57 mutants were screened for degron mutations in all three *AUX/IAA* genes (*IAA1a*/non-annotated, *IAA1b*/*Pp3c8_14720*, and *IAA2/Pp3c24_6610*) and loss-of-function mutations in *DIAGEOTROPICA* (*DGT/ Pp3c1_32280*) by sequencing PCR products. Eighteen new alleles of these loci were identified: *narC32*, *dgt-splice* (g to a 7 bp before ATG); *narD138*, *dgt-S117L*; *narD154*, *dgt-S106C*; *narC15*, *iaa1a-P323L*; *narD69*, *iaa1a-P323L*; *narD75*, *iaa1a-P323L*; *narD148*, *iaa1a-G320S* (weak); *narD151*, *iaa1a-P323L*; *narD153*, *iaa1a-P323L*; *narC33*, *iaa1b-P342F*; *narC37*, *iaa1b-P341L*; *narD68*, *iaa1b-P342L*; *narD152*, *iaa1b-P341L*; *narC8*, *iaa2-P328L*; *narC10*, *iaa2-P328L*; *narC36*, *iaa2-P328F*; *narD139*, *iaa2-P328L*; *narD187*, *iaa2-P327S*. The remaining mutants were compared after growth on BCDAT, BCD, BCD + 0.5 μM NAA, and BCD + 12.5 μM NAA, and twelve were selected based on growth and *DR5* expression phenotypes.

### Sequencing

Genomic DNA was isolated using the *Quick*-DNA Plant/Seed Miniprep Kit (Zymo), and 7.1 to 8.5 total gigabases of paired-end 100 bp reads (BGI Americas) were generated for each mutant plus the *DR5:DsRed*/NLS4 parent strain. Using Bowtie 2 with ‘end-to-end’ and ‘very-sensitive’ settings (v2.3.4.3, ref doi:10.1038/nmeth.1923), the reads were aligned to the Gransden reference sequence to which the *DR5:DsRed* transgene plus scaffold_1157 appended with the correctly assembled *IAA1a* gene were added. The SAM files were converted to sorted BAM files using SAMtools and variants were called using BCFtools (v1.9, ref doi: 10.1093/gigascience/giab008). The variants were mapped relative to the annotations (CoGe GID 33928 v3, Lang et al 2017) using SnpEff (v4.3t, ref: http://dx.doi.org/10.4161/fly.19695).

### Cloning and Molecular Techniques

For ARF cloning, cDNA was synthesized from RNA extraction via RNeasy plant mini kit (Qiagen) and SuperScript III reverse transcriptase (ThermoFisher). Coding sequences were amplified and cloned into pENTR/D-TOPO (ThermoFisher), and subsequently cloned into pGILDA and pB42AD yeast-two-hybrid vectors (Clonetech) via LR Gateway Reactions (ThermoFisher). To introduce T653L and Y643* mutations into ARFd1 entry clone, standard oligomer-driven mutagenesis protocols were used. See primer table for all oligomer sequences.

#### Split YFP

To generate split YFP plasmids, we amplified the N-terminal and C-terminal split YFP cassette described in (Gehl et al., 2009). We cloned each fragment into pMiniT using the PCR Cloning Kit (New England BioLabs), following the manufacturer’s instructions. The resulting plasmids (pMnVN-Gate and pMnVC-Gate) made N-terminal fusions of either the N or C terminal half of mVenus to our protein-of-interest without the excess sequences needed for *Agrobacterium* transformation. Given that each plasmid is Gateway (ThermoFisher) compatible, we were able to use LR reactions with the same entry clones used in Y2H described above. We then transformed 15 μg of each plasmid into *P. patens* protoplasts. These protoplasts were set to incubate in the light for ~60 hours before imaging via confocal microscopy.

### Chlorophyll Extraction

Methanol chlorophyll extractions were performed at room temperature for a minimum of four hours in 2 mL Eppendorf tubes on whole tissue pressed lightly between paper towels to remove excess water. Absorbances were measured on a SmartSpec™Plus spectrophotometer. Chlorophyll content was calculated via the following equation, as described previously(Ritchie, 2006):

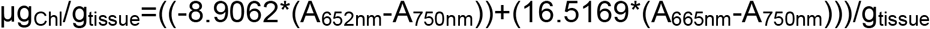

### Image Acquisition, Analysis, and Preparation

For spot-inoculated colony morphology experiments, pictures were acquired via flatbed scanner. Shape parameters were measured in ImageJ based on a binary mask made from color pictures of plants. Images of *arfd1^T653L^* and *arfd1^Y643*^* plants were taken with a Nikon SMZ1500 microscope and a DS-Ri1 camera. For regenerated protoplast assays, images were acquired with BZ-X810 all-in-one microscope (Keyence) using CFP and TRITC filter cubes. Whole plant area was measured in ImageJ using a mask generated from the signal from the calcofluor cell wall stain (Fluorescent Brightener 28, Sigma). Cell length and cross-wall angle were manually measured in ImageJ. Student’s T-Tests were performed in Excel (Microsoft), and the Kolmogorov-Smirnov was done with R. Confocal microscopy was performed on a Zeiss, with image processing and quantification performed in ImageJ.

Dot plots were generated with an internet application (Goedhart, 2021), while histograms and line plots were made with R. Bar plots and histograms were made using Veusz (Qt). Figures were prepared with Affinity Designer (Serif Europe).

### qPCR

cDNA generated from three biological replicates for each condition, with two technical replicates each, were run on a CFX Connect Real Time System thermocycler/plate reader (BioRad). A *P. patens* adenine phosphoribosyltransferase (APT) gene (Pp3c8_16590V3.1) was used as a reference housekeeping gene as described (Bail et al., 2013). See primer table for sequence.

---

**Supplemental Figure 1** Schematic of ARFd mutations, ammonium’s effect on colony morphology A) Models of *PpARFd1* and *PpARFd2*, including where the targets of CRISPR/Cas9-mediated knock out. B) Images of 14-day-old spot inocula on BCD media with or without Ammonium Tartrate (AT). B) Total chlorophyll content via methanol extraction. C) Colony morphology between all lines is largely the same on low-nitogen BCD media.

**Supplemental Figure 2** Schematic of *PpIAA2* and *PpAFB* mutations A) A single nucleotide mutation (red text) in the degron (DII) motif in *PpIAA2* results in a stabilized auxin repressor. B) Alleles recovered when generating *afb1,2,3,4* line. Green text indicates endogenous start codon. Start codon was knocked out in *PpAFB3.* The 2 nt deletion in *PpAFB4* generates a frameshift mutation, resulting in a premature stop codon. The *afb4* peptide is 21 amino acids long.

**Supplemental Figure 3** Schematic of mutagenesis via CRISPR/Cas9 and mutating oligonucleotide. A) Gene diagram of pARFd1::*PpARFd1-mYPet* locus with the PB1 domain in blue. Indicated location of sgRNAs (red text) used to create T653L and Y643* mutations. B) Sequences of WT, *narD122,* mutating oligomer, and recovered *arfd1^T653L^* or *arfd1^Y643*^* mutants. Magenta text highlights codon change, with green highlighting silent mutations intended to introduce restriction enzyme sites. Red/pale red text indicates the protospacer-adjacent motif (PAM), while orange text highlights silent mutations introduced to abolish sgRNA binding. For *arfd1^T653L^,* only the PAM sequence needed to be changed. For *arfd1^Y643*^,* there were no available silent mutations for the lysine residue that would not also disrupt the intron-exon slice site.

**Supplemental Figure 4** Strong auxin signaling mutants exhibit ammonium-dependent caulonemal differentiation A) Representative micrographs of seven-day-old plants regenerated from single protoplasts. WT, *iaa2-mDII,* and *afb1,2,3,4* each exhibit changes in morphology in response to ammonium in the medium. Scale bar is 500 μm. B) Cross wall angles of cells from plants grown on ammonium-deficient medium. Both *iaa2-mDII* and *afb1,2,3,4* protonemata have a smaller cross wall angle compared to wild-type (T-test). This is due to an enrichment of chloronemata (SK test). C) Cross wall angles of cell from plants grown on ammonium-supplemented medium. Each tested line has an increase in chloronemal enrichment, compared to plants grown on ammonium-deficient medium. ** indicates a p-value of <0.001.

**Supplemental Figure 5** Protonemal branching is an auxin-driven developmental processes. A) Protonemal branching occurrence by cell position. Plants were scored after growing for three days on BCD, having regenerated from single protoplasts on PRMB for four days. In addition to two Δ*arfd1,d2* lines, we see a significant delay in the stabilized *iaa2-mDII* line as well as a line lacking any F-box auxin co-receptor, Δ*afb1,2,3,4.* Β) Representative micrographs of protonemal filaments of seven-day-old regenerated protoplasts. White arrowhead indicates filament branch. Scale bar 50 μm.

