## Supplementary figures and images for "A Clade-D Auxin Response Factor is a Major Regulator of Auxin Signaling in *Physcomitrium patens*"

### Supplemental Figure 1

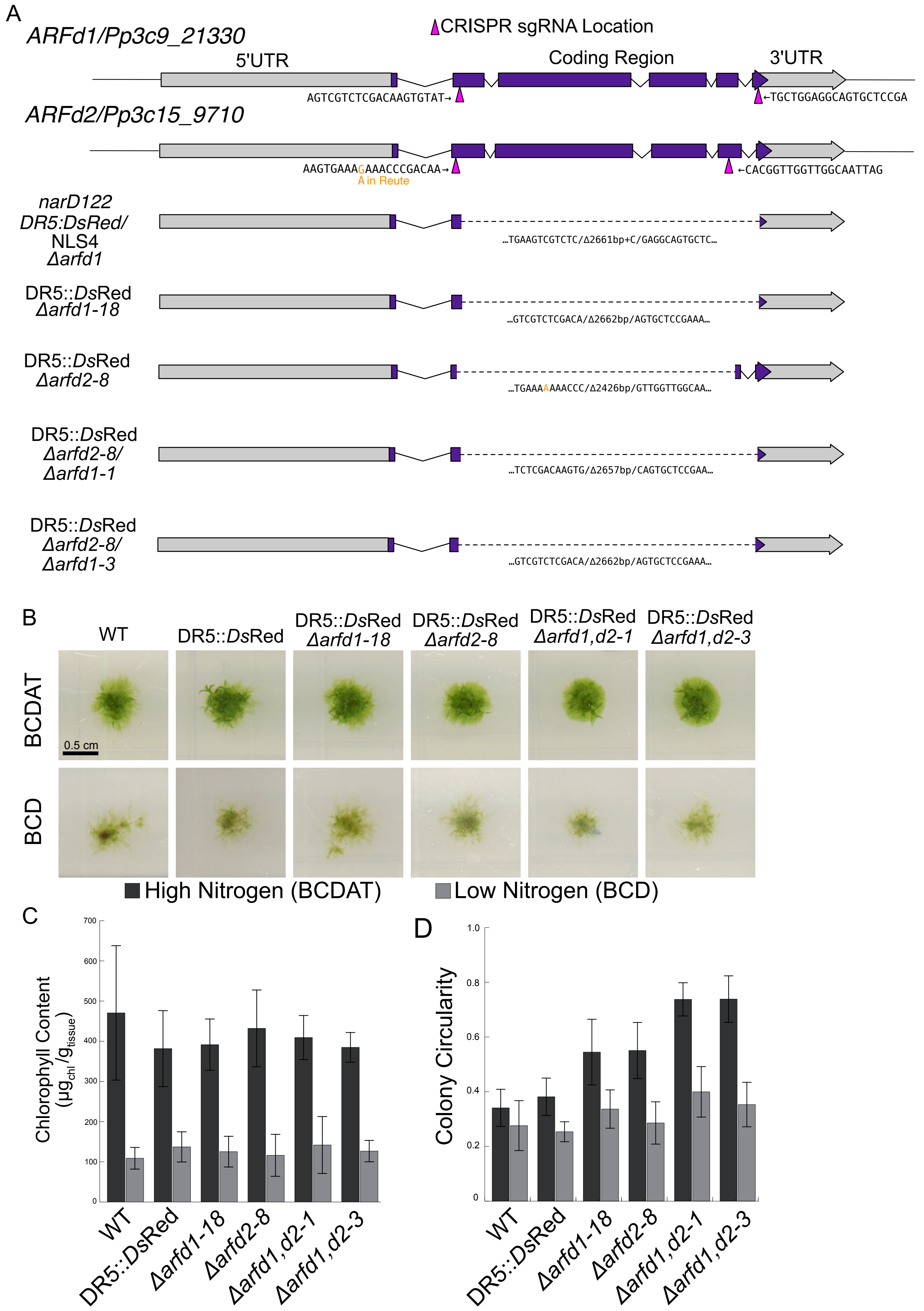

### Supplemental Figure 2

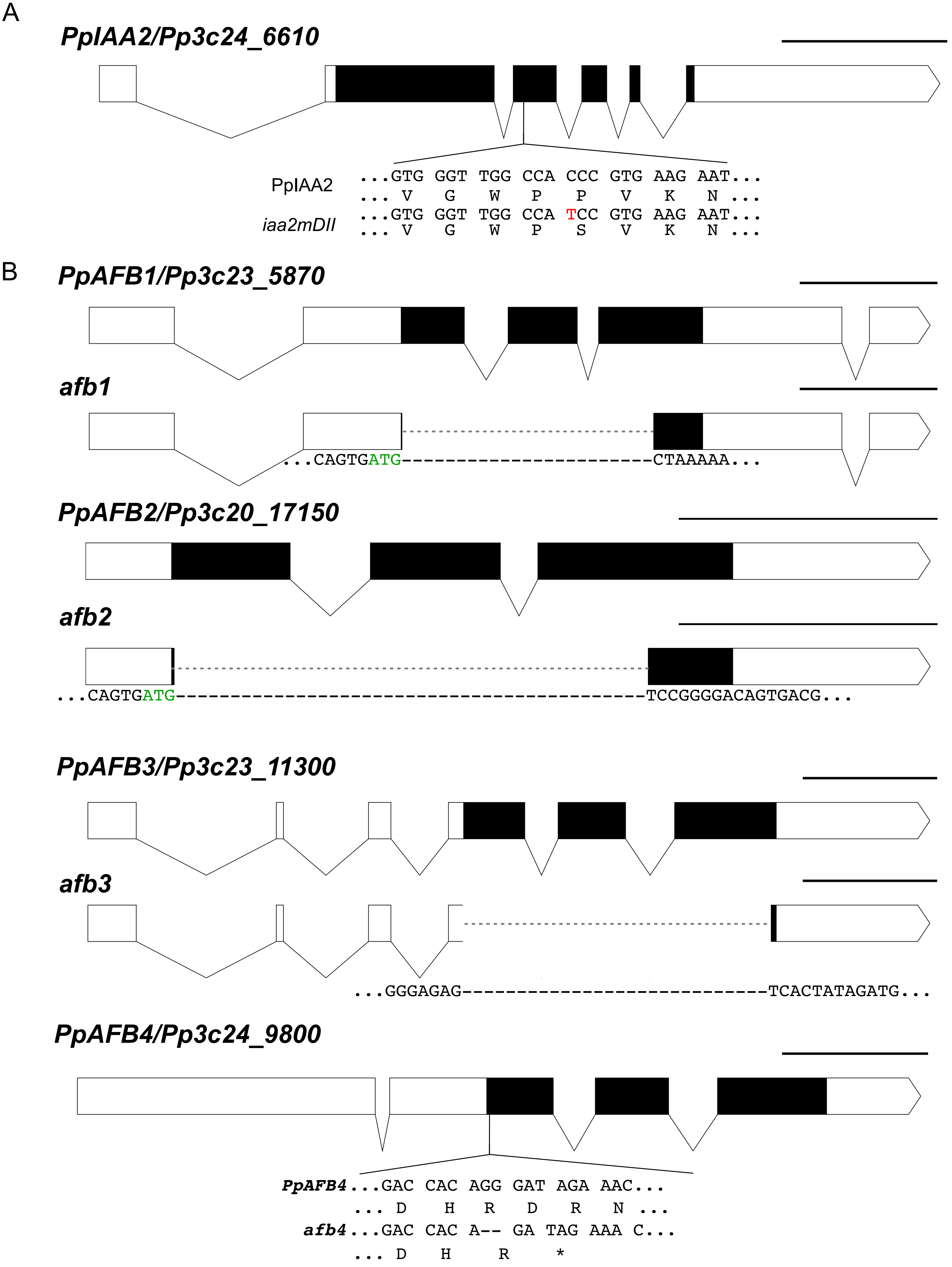

### Supplemental Figure 3

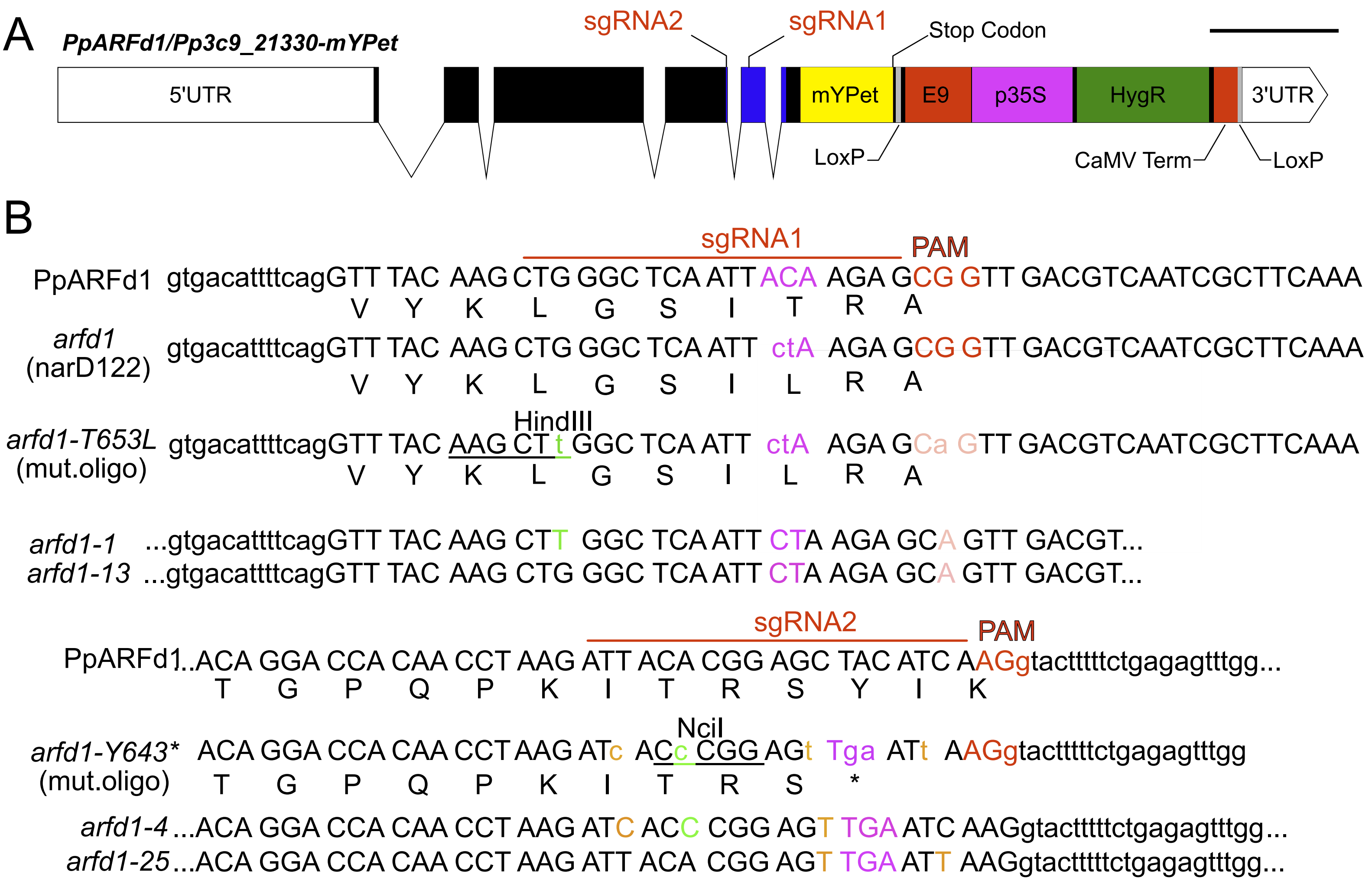

### Supplemental Figure 4

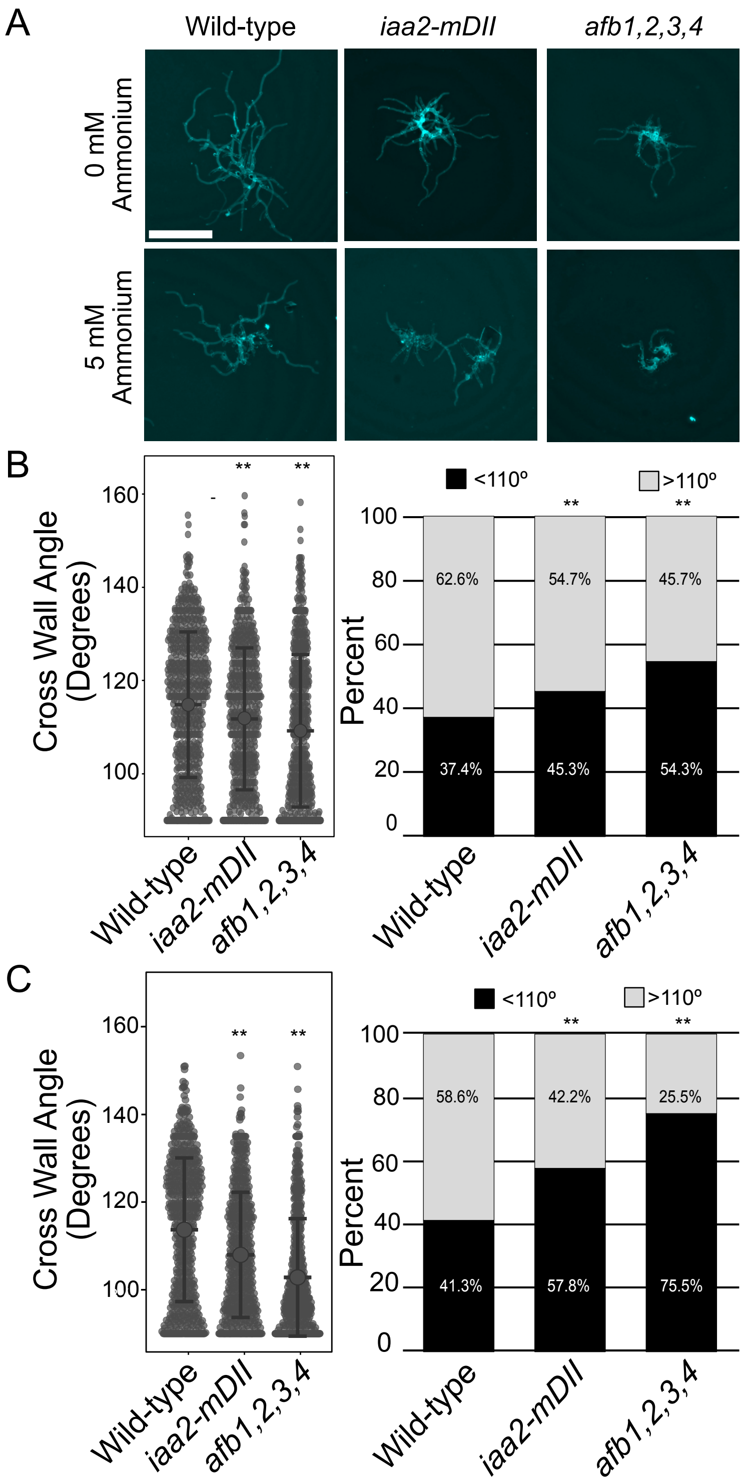

### Supplemental Figure 5

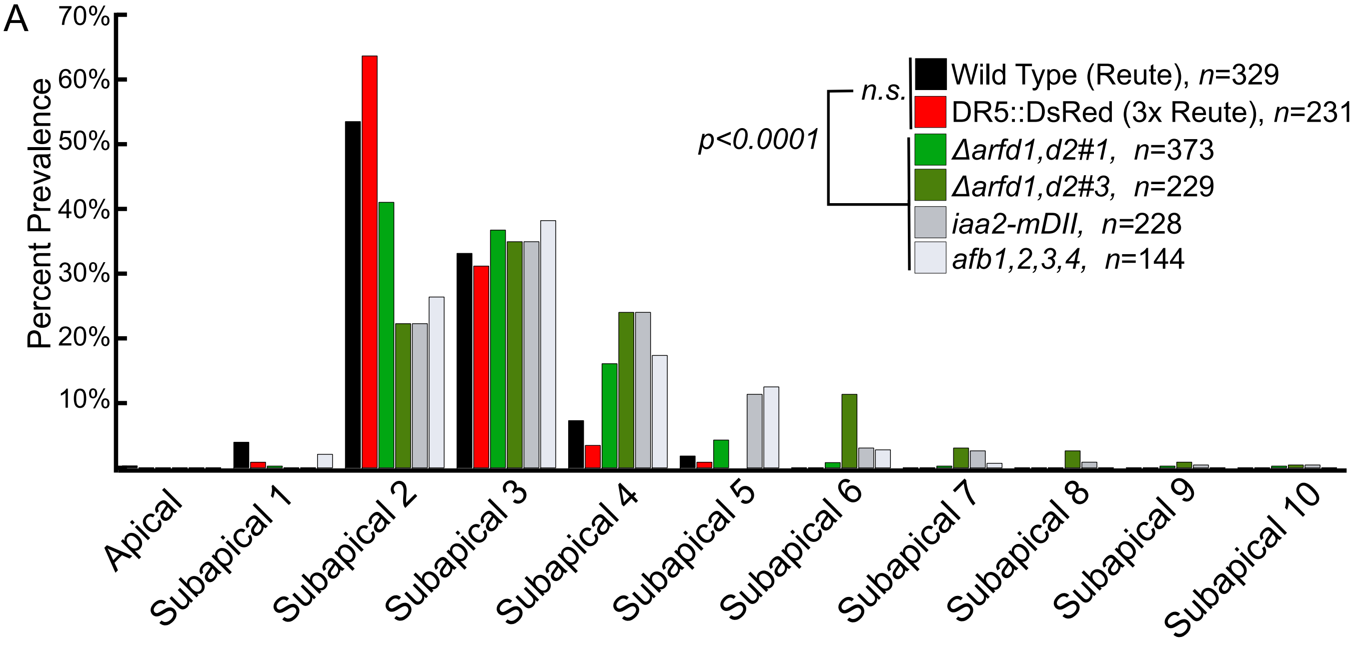

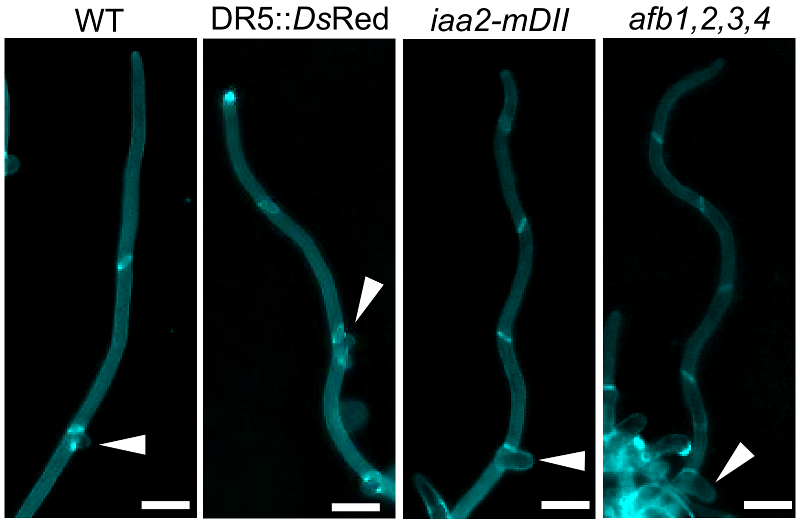
